# YfiM is a lipid-linked oligosaccharide pyrophosphatase

**DOI:** 10.64898/2026.09.16.752124

**Authors:** Sheng-Tao Li, Krzysztof Pawlowski, Victor Lopez, Tatsuya Niwa, Hideki Taguchi, Tadashi Suzuki

## Abstract

Lipid-linked oligosaccharides (LLO), such as Lipid II and O-antigen precursors in bacteria, and dolichol-linked oligosaccharides (DLO) in eukaryotes, are universal metabolic intermediates required for cell wall assembly and protein glycosylation across all domains of life. The oligosaccharide moiety of LLO is synthesized through the sequential addition of various monosaccharides, a process catalyzed by a series of glycosyltransferases, to form the final mature structure. However, the existence of a homeostasis and quality control mechanism to degrade surplus and aberrant LLO has long remained a mystery. Although we recently identified the yeast *LLP1* as an LLO pyrophosphatase involved in the homeostasis and quality control of DLO in fungi, the corresponding mechanism in bacteria and plants has remained elusive. Here, we identify YfiM and AT1G15900 as LLO pyrophosphatases in *Escherichia coli* and *Arabidopsis thaliana*, respectively. Phylogenetic profiling revealed that while YfiM is broadly conserved among Gram-negative bacilli, its plant ortholog displays a unique lineage-specific distribution shaped by selective gene loss. *In vitro*, purified YfiM and AT1G15900 exhibit LLO pyrophosphatase activity. *In vivo*, deletion mutant of *yfiM* caused vancomycin sensitivity in *E. coli*. Conversely, overexpression of YfiM triggered osmotic lysis under hypotonic conditions. Moreover, overexpression of YfiM in an outer membrane-compromised strain partially rescued detergent-chelator sensitivity, revealing a compensatory envelope stress response. Together, our findings uncover an ancient, conserved cross-kingdom system for LLO quality control bridging prokaryotes and plants.

## Introduction

Lipid-linked oligosaccharides (LLO) consist of a hydrophobic lipid tail linked via a high-energy pyrophosphate (PP) bridge to specific carbohydrate chains. They are universal metabolic precursors for protein N-glycosylation and cell envelope assembly across the tree of life (Figure 1) (Aebi, 2013; Manat et al., 2014; Schjoldager et al., 2020). In eukaryotes, the canonical LLO is the dolichol-linked oligosaccharides (DLO), where a Glc_3_Man_9_GlcNAc_2_ core is assembled on a long-chain dolichol pyrophosphate carrier to serve as an essential donor for protein N-glycosylation in the endoplasmic reticulum (Aebi, 2013). On the other hand, prokaryotes predominantly utilize a shorter undecaprenyl lipid carrier (C_55_) to drive the assembly of distinct cell envelope polysaccharides (Kawakami and Fujisaki, 2018). This prokaryotic carrier is dynamically partitioned between synthesizing Lipid II for peptidoglycan cross-linking (Garde et al., 2021) and teichoic acid precursors (Brown et al., 2013) in Gram-positive bacteria, or is diverted toward building lipopolysaccharide (LPS) outer membranes via O-antigen intermediates in Gram-negative bacteria (Bertani and Ruiz, 2018) (Figure 1). Thus, both prokaryotes and eukaryotes rely on LLO as essential precursors for macromolecules critical to cell survival. Consequently, resolving whether species have evolved sophisticated homeostatic and quality control mechanisms to regulate LLO quantity and quality — particularly when subjected to genetic and environmental perturbations — remains a fundamental question in cell biology.

**Figure 1.**
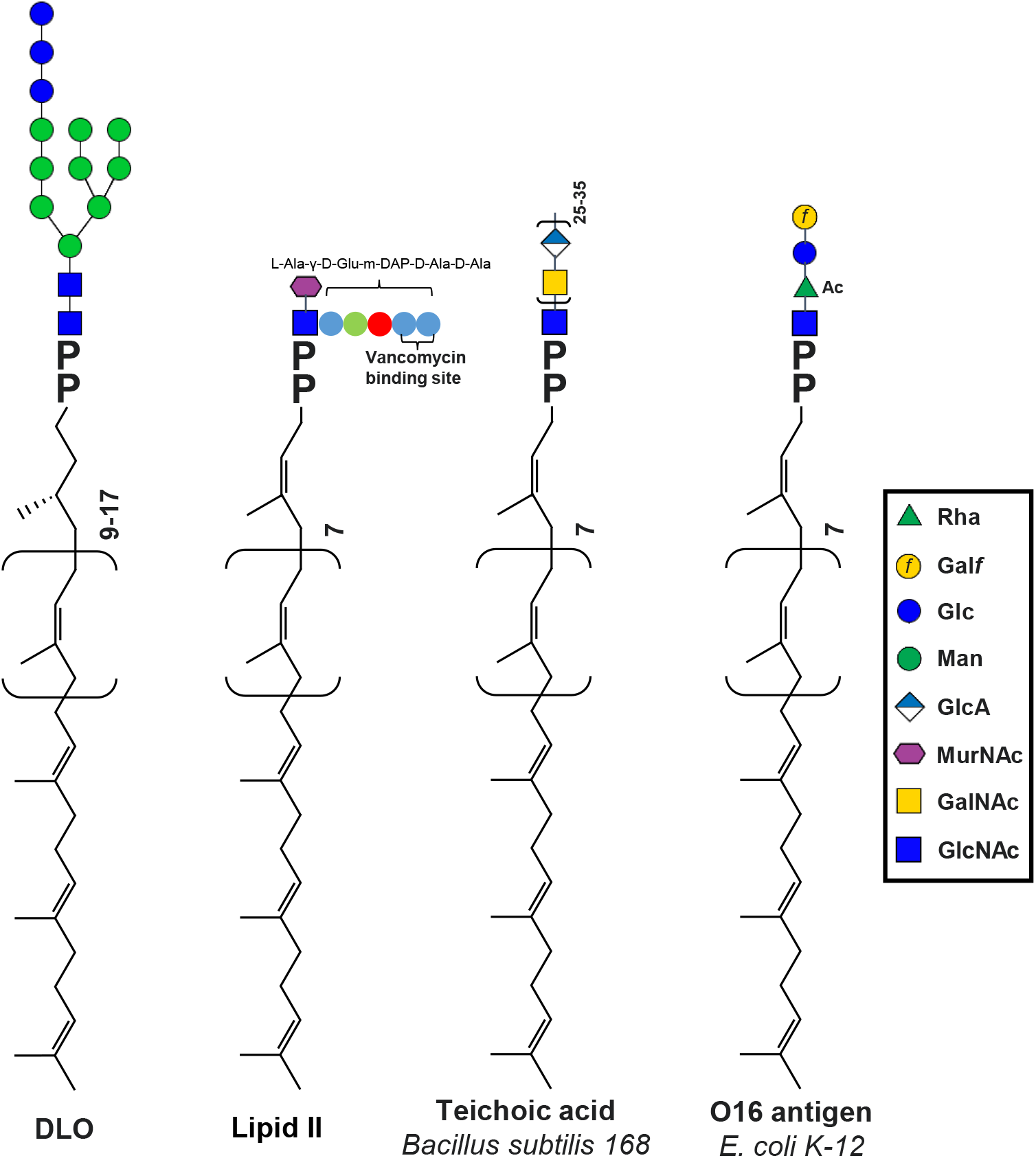
Structural diversity of lipid-linked oligosaccharides (LLO) across domains of life. Schematic representation of the four major LLO intermediates investigated or discussed in this study. All structures share a conserved pyrophosphate (PP) linkage connecting a hydrophobic lipid tail to distinct carbohydrate moieties. From left to right: DLO (dolichol-linked oligosaccharide), the universal precursor for eukaryotic protein N-glycosylation utilizing a dolichol carrier; Lipid II, the conserved peptidoglycan precursor in both Gram-positive and Gram-negative bacteria featuring a short undecaprenyl carrier and an L-Ala-γ-D-Glu-m-DAP-D-Ala-D-Ala pentapeptide; Teichoic acid precursor from *Bacillus subtilis* 168, essential for Gram-positive cell wall functionalization; and O16 antigen precursor from *Escherichia coli* K-12, a representative intermediate for Gram-negative lipopolysaccharide (LPS) outer membrane biogenesis. Monosaccharide symbols are defined in the inset key (Rha, rhamnose; Galf, galactofuranose; Glc, glucose; Man, mannose; GlcA, glucuronic acid; MurNAc, N-acetylmuramic acid; GalNAc, N-acetylgalactosamine; GlcNAc, N-acetylglucosamine; Ac, acetyl group).

We recently identified *Saccharomyces cerevisiae* Llp1 as a novel pyrophosphatase responsible for degrading surplus and aberrant DLO, thereby participating in DLO homeostasis and quality control (Li et al., 2025). Llp1 belongs to the VanZ family proteins. The *vanZ* gene was originally identified in a clinical isolate of *Enterococcus faecium*, BM4147, which is a Gram-positive bacterium, and it confers resistance to the glycopeptide antibiotic vancomycin (Arthur et al., 1995; Arthur et al., 1993; Pootoolal et al., 2002). We previously proposed that VanZ functions as an LLO pyrophosphatase that confers vancomycin resistance by hydrolyzing Lipid II, thereby counteracting vancomycin-induced Lipid II accumulation (Li et al., 2025). However, standard sequence-based homology searches have failed to identify obvious Llp1 or VanZ orthologs in other bacteria, plants or mammals. Given that high-resolution structural prediction tools like AlphaFold can reveal remote structural homologies that evade primary sequence alignments (Abramson et al., 2024), we hypothesized that undiscovered LLO pyrophosphatases — characterized with low sequence identity but conserved tertiary structures and catalytic centers — might exist in these unassigned organisms.

In this study, we identified YfiM, a previously uncharacterized protein in *Escherichia coli*, as a novel LLO pyrophosphatase. While sharing very remote but significant sequence similarity with Llp1 or VanZ, YfiM exhibits striking structural similarity, particularly within its predicted catalytic active site. Utilizing *in vitro* biochemical assays with purified recombinant proteins, we demonstrate that YfiM possesses pyrophosphatase activity toward LLO substrates. Furthermore, phylogenetic analysis of YfiM orthologs revealed that they are ubiquitous across diverse Gram-negative bacteria and widely conserved in the plant kingdom. Together, our findings establish YfiM as a missing link in the evolutionary history of LLO quality control, uncovering a conserved paradigm for lipid carrier recycling and carbohydrate metabolic homeostasis spanning from prokaryotes to plants.

## Results

### Detection of endogenous lipid-linked oligosaccharide pyrophosphatase activity in *Escherichia coli*

Our recent discovery characterized yeast Llp1 as a pyrophosphatase involved in eukaryotic DLO quality control, while its only known bacterial ortholog, VanZ, appears strictly restricted to specific Gram-positive pathogens such as *Enterococcus faecium* (Li et al., 2025). Given the universal requirement for LLO across all cellular life, we hypothesized that analogous LLO pyrophosphatase activities exist in widely distributed Gram-negative bacteria that lack obvious sequence-based orthologs of the Llp1/VanZ family.

To test this hypothesis, we utilized our previously established *in vitro* LLO pyrophosphatase assay. A crude total membrane protein fraction was isolated from the Gram-negative model strain, *E. coli* K-12, as the potential enzyme source (Figure 2A). We then incubated the *E. coli* membrane fraction with Man_5_GlcNAc_2_-DLO as the substrate. To determine whether the lipid carrier was cleaved at the pyrophosphate bridge, the reaction products were treated with or without alkaline phosphatase (AP), subjected to fluorescent labeling (2-aminopyridine), and analyzed via high performance liquid chromatography (HPLC) (Figure 2A), as described previously (Harada et al., 2016). As a result, only completely dephosphorylated glycans are labeled (Hase et al., 1979). Strikingly, a prominent fluorescent glycan peak was detected exclusively in the sample treated with alkaline phosphatase (+AP), whereas no corresponding peak was observed in the untreated control (-AP) (Figure 2B). Thus, the enzymatic product released by *E. coli* membranes is a phosphorylated oligosaccharide. These data directly demonstrate that *E. coli* possess an endogenous biochemical machinery capable of recognizing and hydrolyzing LLO substrates, providing a strong rationale to search for the enzyme responsible for this pyrophosphatase activity.

**Figure 2.**
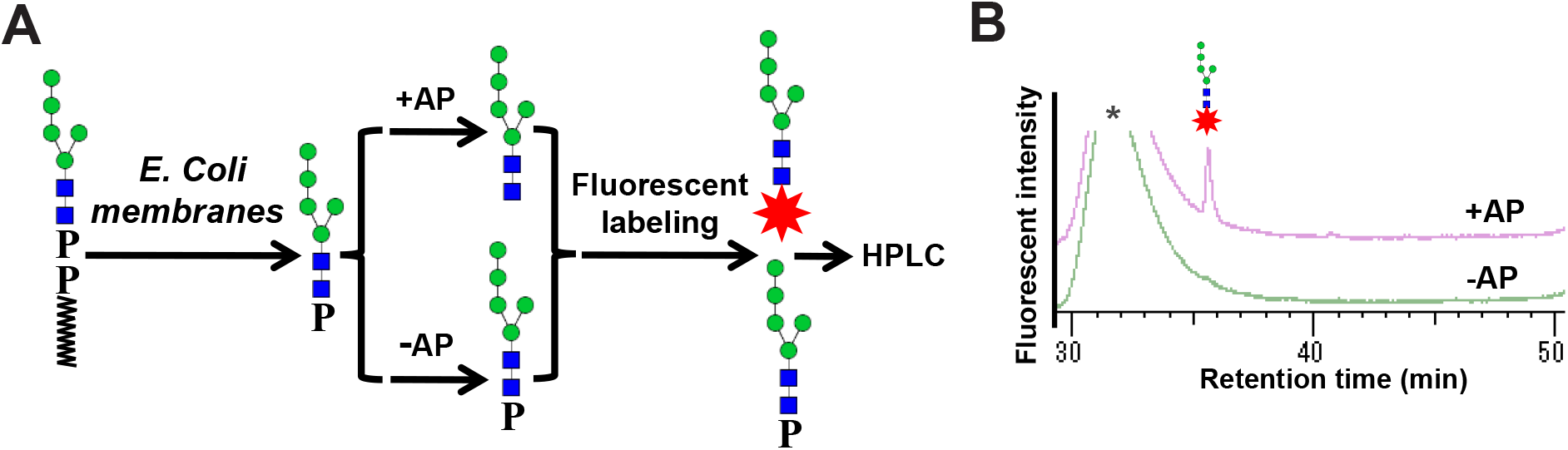
*E. coli* membrane fractions exhibit endogenous LLO pyrophosphatase activity. (A) Schematic workflow of the *in vitro* LLO pyrophosphatase activity assay. DLO substrate was incubated with total membrane proteins isolated from *E. coli* K-12. The reaction product was treated with (+AP) or without (-AP) alkaline phosphatase, followed by fluorescent labeling. The reaction products thus obtained was analyzed by HPLC. (B) HPLC chromatograms of the reaction products. A sharp, fluorescently labeled glycan peak (marked with a red star and schematic glycan structure) eluted at approximately 36 minutes exclusively in the alkaline phosphatase-treated sample (+AP), confirming that the *E. coli* membrane enzyme cleaves the pyrophosphate bond of DLO to release phosphorylated oligosaccharides. An asterisk indicates the nonspecific peak derived from the labeling reagents.

### YfiM shares striking structural similarity and a conserved catalytic center with VanZ

Previous attempts to identify Llp1/VanZ family members by primary sequence in Gram-negative bacteria have been unsuccessful (Li et al., 2025). However, given that enzymes with divergent primary sequences can adopt remarkably similar protein folds to preserve function, we reasoned that a structurally conserved LLO pyrophosphatase might exist in *E. coli*. To uncover remote homologs that evade standard BLAST searches, we performed a Domain Enhanced Lookup Time Accelerated BLAST (DELTA-BLAST) query (Boratyn et al., 2012) against the *E. coli* genome using *Enterococcus faecium* VanZ as the query sequence. This search identified YfiM, an uncharacterized protein annotated as Domain of Unknown Function 2279 (DUF2279), as a potential candidate. Although YfiM shares only ∼16% overall sequence identity and a BLAST E-value of 0.7 with VanZ, a closer examination of the sequence alignment revealed that the identical amino acid residues are clustered around the predicted catalytic active site of VanZ (Figure 3A) (Li et al., 2025). Specifically, the critical residues essential for VanZ activity, including H87, E128, D143, and D147 (Li et al., 2025), align with H31, E72, D87, and D91 of YfiM, respectively (Figure 3A).

**Figure 3.**
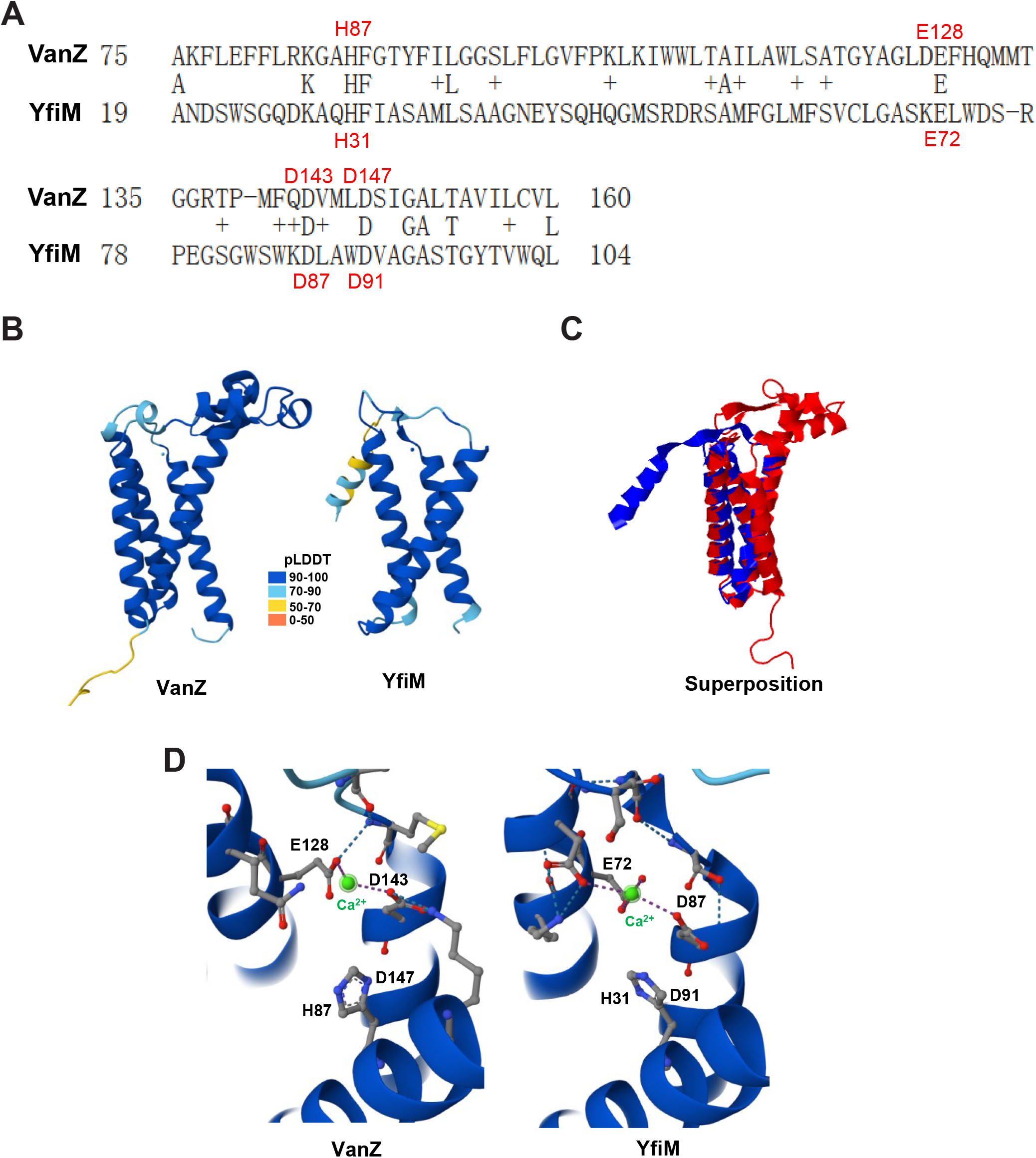
Bioinformatic and structural alignment reveals YfiM as a sequence divergent, yet structurally conserved homolog of VanZ. (A) DELTA-BLAST sequence alignment between *Enterococcus faecium* VanZ and *E. coli* YfiM. Conserved and identical residues are highlighted, revealing a strict alignment of key catalytic residues (VanZ H87, E128, D143, D147 correspond to YfiM H31, E72, D87, D91). (B) Overall tertiary structural comparison of AlphaFold predicted models for VanZ (left) and YfiM (right), colored by the per-residue confidence metric pLDDT on a scale from 0 to 100. (C) Superposition of VanZ (red) and YfiM (blue). (D) Close up view of the highly conserved catalytic center.

To gain insight into this divergent homolog, we compared the AlphaFold structures of VanZ and YfiM (Abramson et al., 2024). VanZ is predicted to possess four transmembrane domains (TMDs), whereas YfiM contains only three (Figure 3B).

However, analysis using TM-align for their protein structure alignment and comparison yielded an RMSD of 2.3 and a TM-score of 0.7, indicating that they share the same structural fold (Figure 3C) (Zhang and Skolnick, 2005). Importantly, alignment of their predicted active sites revealed that the spatial coordination of the residues required for Ca^2+^ coordination was virtually identical (Figure 3D). It is worth noting that recent bioinformatics analyses suggest VanZ and YfiM belong to the lipocone superfamily and are involved in the metabolism of lipids, peptidoglycan, and exopolysaccharides (Burroughs et al., 2025). Taken together, these data suggest that YfiM is a member of the Llp1/VanZ pyrophosphatase family in *E. coli*.

### YfiM possesses LLO pyrophosphatase activity and is broadly conserved across gram-negative bacteria

To experimentally validate whether YfiM is indeed the enzyme responsible for the LLO pyrophosphatase activity, we purified recombinant *E. coli* YfiM and mutants that were predicted to be catalytically inactive. YfiM and the mutants were expressed and purified as fusion proteins with a Mistic tag on their N-terminus (Figure 4A) (Roosild et al., 2005). To test the catalytic activity of YfiM we utilized our established *in vitro* assay. Indeed, purified Mistic-YfiM but not Mistic tag alone exhibited pyrophosphatase activity (Figure 4B). The predicted active site of YfiM contains a putative metal-binding pocket, therefore, we examined its divalent cation dependence. YfiM activity was strictly dependent on exogenous divalent metals; it exhibited maximum catalytic efficiency in the presence of Ca^2+^, partial activity with Mn^2+^, and was completely inactive when treated with Co^2+^, Mg^2+^, or the metal chelator EDTA (Figure 4C), which is consistent with Llp1/VanZ (Li et al., 2025). To test the predicted active site observed in the AlphaFold model, we evaluated the catalytic activity of several mutants (K28, H31, E72, and D91). Remarkably, alanine substitution of any of these residues led to a near complete or total loss of LLO pyrophosphatase activity (Figure 4D). These results collectively demonstrate that YfiM is a LLO pyrophosphatase *in vitro*.

**Figure 4.**
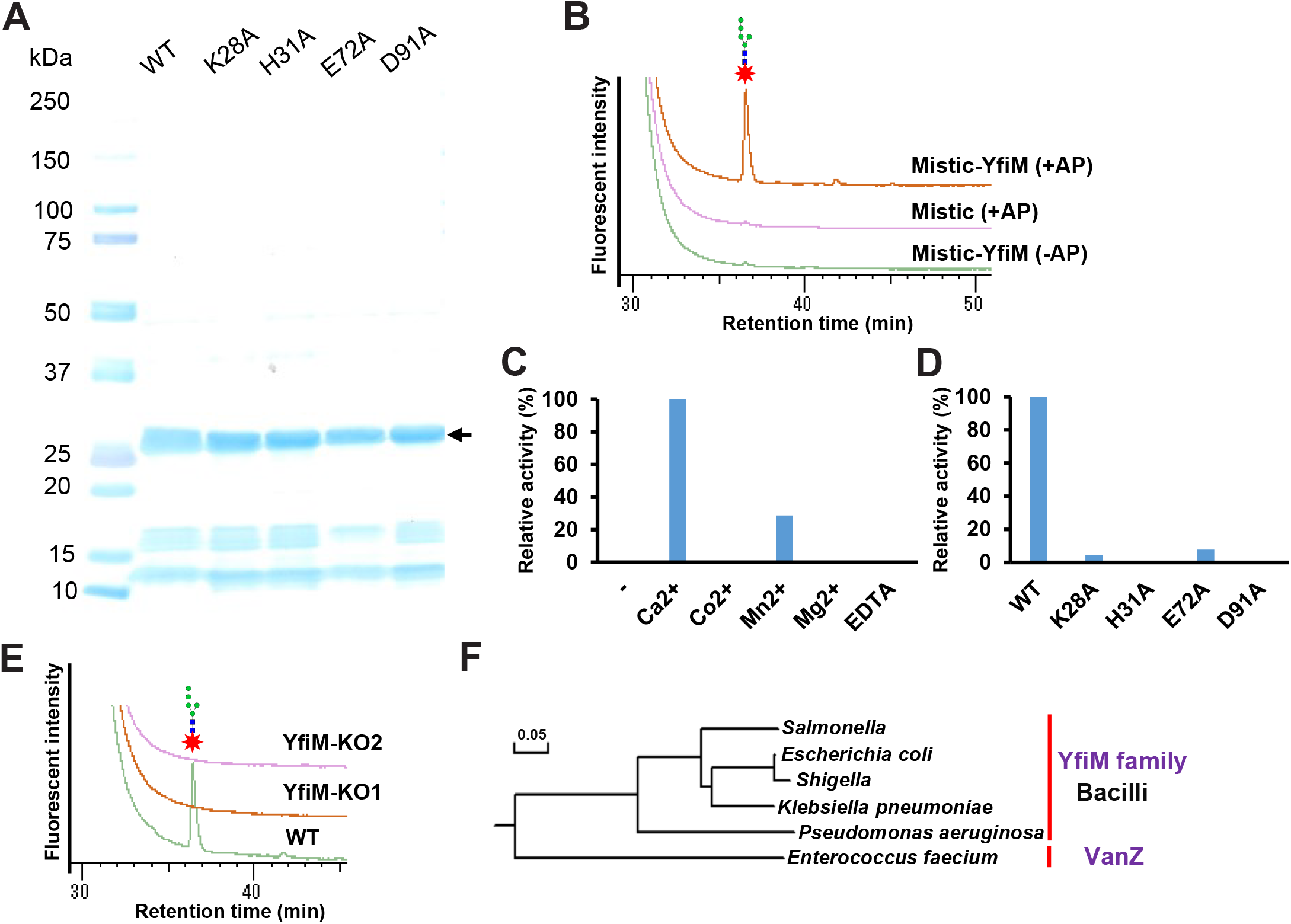
Biochemical characterization, mutational analysis, and phylogenetic distribution of YfiM. (A) SDS-PAGE analysis of purified recombinant wild type (WT) Mistic-YfiM and its catalytic site mutants (K28A, H31A, E72A, D91A). Molecular weight markers (kDa) are indicated on the left. The arrow indicated Mistic-YfiM. (B) HPLC profiles confirming the *in vitro* LLO pyrophosphatase activity of purified Mistic-YfiM using Man_5_GlcNAc_2_-DLO as a substrate. (C) Divalent cation dependence of YfiM. Relative pyrophosphatase activity was measured in the presence of various metal ions (Ca^2+^, Co^2+^, Mn^2+^, Mg^2+^) or EDTA. Activity measured in the presence of Ca^2+^ was set to 100%. (D) Relative enzymatic activity of WT Mistic-YfiM and the indicated alanine mutants, demonstrating the indispensability of the conserved pocket residues. (E) HPLC profiles obtained from the LLO pyrophosphatase assay using total membrane proteins isolated from *E. coli* WT or two independent *yfiM* knockout strains. (F) Phylogenetic tree of the YfiM family. YfiM orthologs form a tightly clustered clade strictly conserved among Gram-negative bacilli (rod-shaped bacteria, including *Salmonella, Escherichia coli, Shigella, Klebsiella pneumoniae*, and *Pseudomonas aeruginosa*). This family stands evolutionary segregated from the Gram-positive *Enterococcus faecium* VanZ clade (scale bar indicates genetic distance).

To investigate the physiological relevance of YfiM, we next examined the LLO pyrophosphatase activity in *E. coli* gene deletion strains (Baba et al., 2006; Yamamoto et al., 2009). Strikingly, while total membrane fractions isolated from the wild type (WT) strain exhibited robust activity, this activity was completely abolished in two independent *yfiM* knockout strains (Figure 4E). This demonstrates that YfiM is the sole, non-redundant enzyme responsible for endogenous LLO pyrophosphatase activity in *E. coli*.

Given the distant sequence similarity of YfiM, we set out to explore the evolutionary distribution of this newly identified pyrophosphatase. We conducted a comprehensive phylogenetic analysis across diverse Gram-negative bacteria, which are broadly categorized into cocci, spiral, and bacilli forms based on cell morphology (Nagaraja, 2022; Pavlova et al., 2022). Remarkably, our analysis revealed that YfiM is not universally distributed but is strictly and highly conserved within the bacilli lineage (Figure 4F). The phylogenetic tree demonstrates that YfiM orthologs form a distinct, coherent evolutionary clade comprising major Gram-negative rod-shaped pathogens, including Salmonella, Shigella, Klebsiella, and Pseudomonas (Figure 4F). Conversely, no obvious YfiM counterparts were identified in Gram-negative cocci or spiral bacteria. This narrow distribution within the bacilli group stands evolutionary segregated from the Gram-positive *Enterococcus faecium* VanZ clade, highlighting a distinct evolutionary adaptation tailored to the cell envelope homeostasis of Gram-negative bacilli. Taken together, these data establish YfiM as the definitive, metal-dependent LLO pyrophosphatase in Gram-negative bacteria.

### Deletion or overexpression of YfiM alters osmotic stability and outer membrane permeability barriers

To elucidate the physiological function of YfiM and its impact on the *E. coli* cell envelope, we characterized the phenotypic consequences of both *yfiM* deletion and overexpression. Given that VanZ confers resistance to glycopeptide antibiotics by modulating lipid II availability (Arthur et al., 1995; Li et al., 2025; Pootoolal et al., 2002), we first investigated whether YfiM exerts a similar protective effect against vancomycin in *E. coli* cells. Under normal conditions, the Gram-negative outer membrane acts as an intrinsic permeability barrier that restricts vancomycin influx due to its large molecular weight (Nikaido, 2003; Silhavy et al., 2010). Notably, the *yfiM* knockout strain exhibited significantly increased sensitivity to vancomycin compared to the parental wild-type strain (BW25113-WT) (Figure 5A). On the other hand, overexpression of YfiM in the *E. coli* Rosetta strain (pET28-YfiM) conferred somewhat enhanced resistance to vancomycin relative to the empty vector control (Rosetta-pET28) (Figure 5A). These findings demonstrate that YfiM can modulate cellular sensitivity to cell wall targeting antibiotics *in vivo*, reminiscent of the VanZ family.

**Figure 5.**
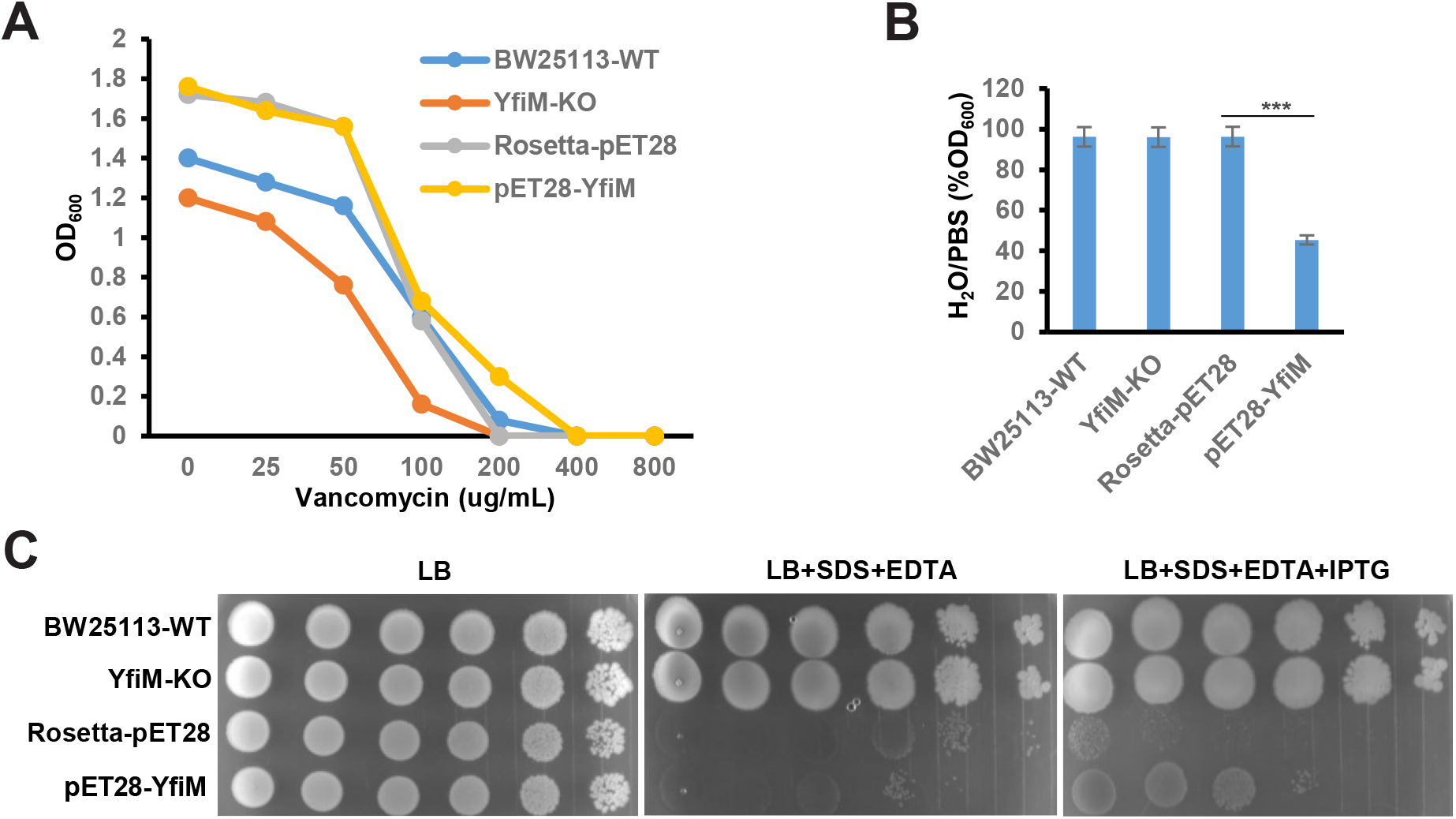
Phenotypic consequences of *yfiM* disruption or overexpression on antibiotic resistance, osmotic stability, and cell envelope integrity. (A) Vancomycin susceptibility curves of *E. coli* strains. Cell density (OD_600_) was measured following incubation with indicated concentrations of vancomycin (0-800 μg/mL). (B) Osmotic shock assay assessing peptidoglycan structural integrity. Cells were resuspended in either isotonic PBS or pure water, and the ratio of residual optical density (H_2_O/PBS, % OD_600_) was calculated. Statistical significance was determined using Student’s t-test (***, p < 0.001). Data represent mean ±SD from three independent biological replicates. (C) Serial 10-fold dilution spotting assays on LB agar plates under the indicated stress conditions (LB only, LB + SDS + EDTA, or LB + SDS + EDTA + IPTG).

We hypothesized that the peptidoglycan intermediate, lipid II, is a primary substrate for YfiM. Thus, we reasoned that uncontrolled overexpression of this pyrophosphatase would deplete this critical precursor, thereby compromising the structural integrity of the peptidoglycan layer (Manat et al., 2014). Given that the peptidoglycan sacculus acts as the primary mechanical determinant for maintaining internal turgor pressure and osmotic stability in bacteria (Garde et al., 2021), we subjected cells to severe osmotic shock by shifting them from an isotonic buffer (PBS) to pure water, monitoring cell lysis via the residual OD_600_ ratio (H_2_O/PBS). While the WT, *yfiM*-KO, and empty vector control strains remained fully intact upon water resuspension, YfiM overexpressing cells (pET28-YfiM) experienced catastrophic osmotic lysis, retaining only ∼40% of their initial optical density (Figure 5B). This osmotic fragility strongly indicates that overexpression of YfiM causes severe, uncontrolled degradation of lipid-linked cell wall precursors, leading to a structurally defective peptidoglycan meshwork.

Finally, we evaluated the impact of YfiM perturbation on outer membrane (OM) integrity by measuring sensitivity to the anionic detergent SDS in the presence of EDTA, a divalent cation chelator that disrupts the LPS interaction network (Guest et al., 2023; Leive, 1965; Nikaido, 2003). In serial dilution spotting assays, the BW25113-WT and *yfiM*-KO strains, which are both derivatives of *E. coli* K-12, displayed robust growth on plates containing SDS and EDTA (Figure 5C). Although K-12 strains harbor a mutation disrupting O-antigen, they retain a complete and structurally intact LPS core oligosaccharide (Jeong et al., 2009; Studier et al., 2009), which provides sufficient barrier function against detergent entry (Nikaido, 2003). Conversely, the Rosetta strain (a derivative of *E. coli* B) possesses an intrinsically truncated core oligosaccharide background (Jeong et al., 2009; Studier et al., 2009), rendering it hypersensitive to SDS-EDTA (Figure 5C). Strikingly, upon induction with IPTG, overexpression of YfiM in the Rosetta background (pET28-YfiM) partially restored growth and rescued the strain from SDS-EDTA toxicity (Figure 5C). We hypothesize that this counterintuitive resistance arises from a cellular compensatory mechanism: the severe peptidoglycan damage triggered by YfiM overexpression likely elicits a robust envelope stress response which prompts the bacteria to structurally compensate by increasing outer membrane thickness, density, or lipoprotein cross-linking to ensure survival under stress (Grabowicz and Silhavy, 2017; Saha et al., 2021). Together, these genetic and phenotypic analyses suggest that YfiM is involved in governing cell envelope homeostasis and mechanical strength in *E. coli*.

### Identification of DLO pyrophosphatase activity of plant YfiM orthologs

The strict restriction of YfiM to Gram-negative bacilli, combined with its role in bacterial cell envelope homeostasis, prompted us to explore whether YfiM was horizontally transferred or selectively inherited across major biological kingdoms during ancient evolutionary history. Remarkably, a standard primary sequence alignment by BLAST using YfiM as a direct query sequence successfully identified uncharacterized, sequence-conserved orthologs within the plant kingdom, with a representative candidate gene found in the model plant *Arabidopsis thaliana*, designated as *AT1G15900* (Figure 6A). To map its evolutionary trajectory, we performed a comprehensive phylogenetic analysis across diverse photosynthetic lineages. This analysis indicated a unique, non-contiguous distribution, with *AT1G15900* homologs found in algae, mosses, and angiosperms, but absent in monilophytes (ferns) and gymnosperms (Figure 6B). This disparate pattern highlights a potential case of selective, lineage-specific gene loss during land plant evolution — a phenomenon well-documented in plant comparative genomics where non-essential or functionally redundant pathways are purged in specific lineages while being strictly retained in others (Albalat and Cañestro, 2016; Clark, 2023). Despite this evolutionary mosaic, the sequence conservation between Gram-negative bacilli and both lower and higher plants underscores an ancient, foundational role for this pyrophosphatase.

**Figure 6.**
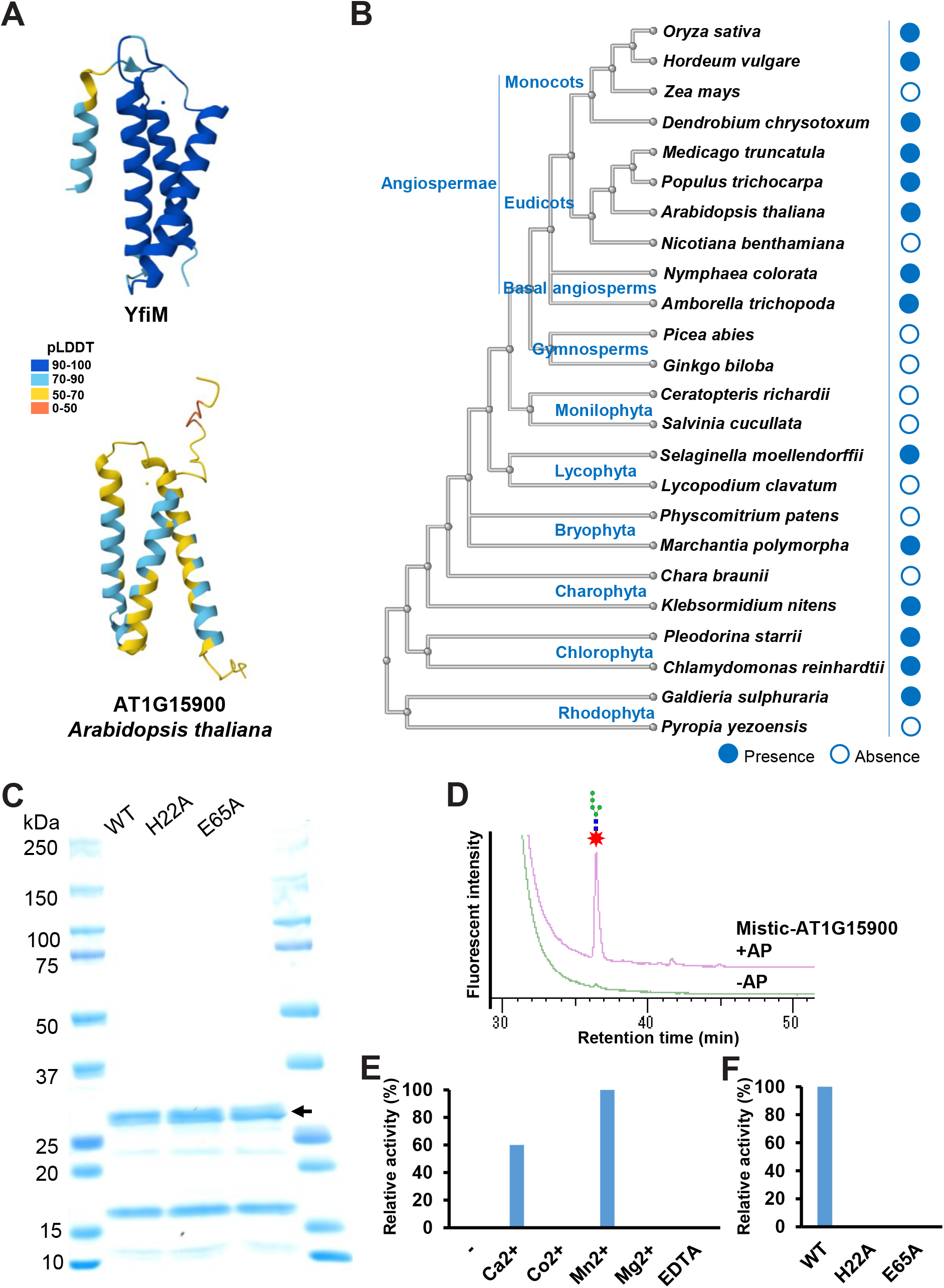
Structural conservation, phylogenetic distribution, and biochemical characterization of the plant YfiM ortholog AT1G15900. (A) Three-dimensional structural comparison of AlphaFold predicted models for *E. coli* YfiM (top) and *Arabidopsis thaliana* AT1G15900 (bottom), colored by the per-residue confidence metric pLDDT on a scale from 0 to 100. (B) Phylogenetic tree illustrating the evolutionary distribution of AT1G15900 orthologs across the plant kingdom. Filled blue circles indicate presence, and open circles indicate absence. (C) SDS-PAGE analysis of purified recombinant WT Mistic-AT1G15900 and its predicted active site mutants (H22A and E65A). Molecular weight markers (kDa) are indicated on the left. The arrow indicated Mistic-AT1G15900. (D) HPLC profiles of *in vitro* pyrophosphatase assays using purified Mistic-AT1G15900. (E) Divalent cation dependence of AT1G15900. Relative pyrophosphatase activity was measured in the presence of various metal ions or EDTA. Activity measured in the presence of Mn^2+^ was set to 100%. (F) Relative enzymatic activity of WT Mistic-AT1G15900 and the H22A and E65A mutants.

To experimentally determine whether this plant ortholog functions as a *bona fide* LLO (DLO)-pyrophosphatase, we expressed and purified AT1G15900 and its active site mutants as Mistic fusion proteins (Figure 6C). Utilizing our *in vitro* assay, purified Mistic-AT1G15900 exhibited robust DLO pyrophosphatase activity (Figure 6D). We next analyzed the biochemical properties and catalytic requirements of AT1G15900. Like its bacterial counterpart, AT1G15900 activity was strictly dependent on exogenous divalent cations (Ca^2+^ or Mn^2+^) (Figure 6E). Furthermore, we mutated the invariant residues predicted to coordinate cation to alanine (H22A and E65A). Indeed, both the H22A and E65A substitutions completely abolished the LLO pyrophosphatase activity of AT1G15900 (Figure 6F). Taken together, these results confirm that AT1G15900 has LLO pyrophosphatase activity, revealing a mechanism of lipid carrier recycling that is highly conserved from prokaryotes to plants.

## Discussion

Although DLO pyrophosphatase activity has been known in mammals and yeast since the 1970’s (Cacan et al., 1980; Cacan et al., 1989; Harada et al., 2013b; Hsu et al., 1974; Massarweh et al., 2016), the underlying gene remained a mystery until our recent identification of *LLP1* in yeast (Li et al., 2025). Because Llp1 and its homolog VanZ were restricted to fungi and *Enterococcus faecium*, our knowledge of the evolutionary distribution of this enzyme remained limited. Here, we expand these boundaries by identifying YfiM and its orthologs as definitive LLO pyrophosphatases in Gram-negative bacteria and plants. Our findings reveal a structurally and mechanistically conserved, cross-kingdom enzymatic paradigm that governs glycosyl carrier lipid salvage across the domains of life.

The multilayered Gram-negative cell envelope, which includes the inner membrane, the peptidoglycan-containing periplasm, and the asymmetric outer membrane, acts as a formidable barrier protecting bacteria from environmental stressors, host immunity, and antimicrobials (Nikaido, 2003; Silhavy et al., 2010). The biogenesis of this complex architecture heavily relies on LLO, which serve as essential biosynthetic intermediates for the peptidoglycan precursor Lipid II (Garde et al., 2021) and the lipopolysaccharide O-antigen (Bertani and Ruiz, 2018). Crucially, in the *Enterobacteriaceae* family such as *E. coli*, this LLO dependency extends to other major cell surface polysaccharides, including colanic acid (Scott et al., 2019) and the enterobacterial common antigen (ECA) (Bennett and Mitchell, 2025). The *de novo* synthesis of their shared glycosyl carrier lipid, undecaprenyl phosphate (C_55_-P), is an energetically expensive process. Thus, cells maintain a strictly limited intracellular pool of C_55_-P that must be continuously recycled (Manat et al., 2014). Consequently, perturbing any single polysaccharide assembly pathway, whether genetically or chemically, can cause intermediate accumulation, trapping the finite carrier pool in a metabolic dead-end. This could then initiate a catastrophic domino effect that cripples the synthesis of all other surface polysaccharides. Whether YfiM operates as a salvage factor (functionally analogous to yeast Llp1 in maintaining DLO homeostasis and quality control) represents a critical future avenue to dissect the quality control networks governing the Gram-negative cell envelope. Concurrently, investigating whether the plant ortholog, AT1G15900, plays a parallel regulatory role in gating the dolichol phosphate (Dol-P) pool to modulate eukaryotic DLO homeostasis and protein N-glycosylation will be vital to fully comprehend the physiological significance of this ancient pyrophosphatase family across kingdoms (Strasser, 2016).

Given the strict restriction of VanZ to specific Gram-positive *Enterococcus faecium* and YfiM to Gram-negative bacilli, we sought to determine whether LLO quality control machinery is universally integrated into the cell envelope biogenesis of other divergent bacterial lineages. By leveraging DELTA-BLAST queries across a diverse panel of model bacterial genomes, we identified structural and catalytic-site conserved orthologs across distinct phyla, which we classified into Type 1 and Type 2 groups based on their distinct transmembrane helical topologies (Figures S1A-E). Despite sharing conserved active sites with VanZ, these recombinant Type 1 and Type 2 enzymes failed to exhibit detectable DLO pyrophosphatase activity in our *in vitro* assays using the eukaryotic DLO substrate. Whether these uncharacterized proteins possess LLO pyrophosphatase activity with narrow substrate specificity requires future testing utilizing native bacterial LLO substrates.

Ultimately, through our present and recent work, LLO pyrophosphatase genes have now been successfully defined across prokaryotes and the plant kingdom, in addition to yeast (Li et al., 2025). However, the molecular identity of the mammalian LLO pyrophosphatase has eluded discovery for decades. Intrigued by this phylogenetic gap, our comprehensive genomic database queries failed to detect any obvious structural or catalytic-site orthologs of either the Llp1/VanZ or YfiM lineages in vertebrate genomes. This structural divergence is perhaps unsurprising given the profound differences in their biochemical properties. While the bacterial and plant enzymes characterized herein are driven by Ca^2+^, *in vitro* studies reported that the endogenous mammalian LLO pyrophosphatase activity is dependent on Co^2+^ and entirely inert in the presence of Ca^2+^ (Harada et al., 2016; Li et al., 2025; Massarweh et al., 2016). This strongly implies that mammals have evolved a structurally disparate and mechanistically distinct enzyme to manage LLO turnover. Consequently, unmasking the mammalian LLO pyrophosphatase will be an indispensable milestone in fully comprehending the diverse physiological underpinnings of carbohydrate metabolic homeostasis across the entire tree of life.

## Materials and Methods

### Bacterial strains and culture

*E. coli* wild-type K-12 strain BW25113 was kindly provided by Dr. Shintaro Iwasaki (RIKEN Pioneering Research Institute). The *E. coli yfiM* knockout strains (two clones) in the BW25113 background were derived from the Keio collection (Baba et al., 2006; Yamamoto et al., 2009). The *E. coli* B Rosetta 2 (DE3) strain was purchased from Sigma-Aldrich (71401). *E. coli* was cultured in LB medium (1% tryptone, 0.5% yeast extract, and 1% NaCl) or TB medium (1.2% tryptone, 2.4% yeast extract, 0.5% glycerol, 17 mM KH_2_PO_4_, and 72 mM K_2_HPO_4_) at 37°C. The media were supplemented with the following as needed: 50 μg/mL kanamycin, 34 μg/mL chloramphenicol, 25-800 μg/mL vancomycin, 0.5% SDS, 0.5 mM EDTA, and 0.2 mM IPTG.

### Plasmid construction

YfiM and the N-terminally Mistic-tagged YfiM were constructed in the pET28 vector to generate pET28-YfiM and pET28-Mistic-YfiM, respectively. N-terminally Mistic-tagged AT1G15900 were constructed in the pET28 vector to generate pET28-Mistic-AT1G15900. Various point mutants were generated using QuikChange Lightning Site-Directed Mutagenesis Kit (210518; Agilent Technologies). The sequence of the plasmid was confirmed by direct DNA sequencing.

### Bioinformatic analyses

To build the YfiM phylogenetic tree, we searched for homologs of *E. coli* YfiM in a curated database containing the proteomes of *Salmonella, Shigella, Klebsiella pneumoniae*, and *Pseudomonas aeruginosa* using the NCBI BLAST program blastp. Protein sequences of the YfiM homologs and VanZ were aligned using DNAMAN software tool. A phylogenetic tree was subsequently constructed from the aligned sequences using the Observed Divergency method within DNAMAN. The same method was used to construct the phylogenetic trees for VanZ type 1 and type 2.

To build the phylogenetic tree illustrating the evolutionary distribution of AT1G15900 orthologs across the plant kingdom, we searched for homologs of AT1G15900 in a total of 24 plant species with high-quality genomes from representative taxonomic groups (Rhodophyta, Chlorophyta, Charophyta, Bryophyta, Lycophyta, Monilophyta, Gymnosperms, Basal angiosperms, Eudicots and Monocots) using the NCBI BLAST program blastp. The phylogenetic tree of the studied 24 plant species was constructed using the NCBI CommonTree tool based on the NCBI Taxonomy database (Schoch et al., 2020). The lineage data for all target taxa were retrieved and compared to build the hierarchical clustering structure reflecting their evolutionary relationships. The resulting tree file was subsequently visualized and formatted using NCBI Tree Viewer.

### Preparation of membrane fractions from *E. coli* K-12 BW25113

A 1 mL saturated *E. coli* K-12 BW25113 wild type or *yfiM* knockout strain overnight culture in LB medium was inoculated into a 50 mL LB medium and grown for another 5 h. The medium was removed by centrifugation. The cell pellet was washed twice with 40 mL buffer A (20 mM Tris-HCl, pH 7.5, 150 mM NaCl) and resuspended in 10 mL buffer A, then disrupted by sonication. Cell lysates were centrifuged at 5000 × *g* for 5 min at 4°C to remove cellular debris. Membrane fractions were collected by centrifugation at 20,000 × *g* for 1.5 h at 4°C and resuspended in 0.5 mL buffer A for the LLO pyrophosphatase assay.

### Expression and purification of recombinant YfiM and AT1G15900 from *E. coli*

To purify recombinant Mistic-YfiM (WP_001300818.1) and Mistic-AT1G15900 (NP_173042.1), pET28-Mistic-YfiM or pET28-Mistic-AT1G15900 were transformed into *E. coli* Rosetta 2 (DE3) cells, and positive clones were selected by kanamycin and chloramphenicol. Single colonies were cultured in TB medium containing kanamycin and chloramphenicol at 37°C until OD_600_ was between 0.8 and 1.2. Cultures were cooled to 16°C, and protein expression was induced with 0.2 mM IPTG. After 16 h, cells were collected, resuspended in buffer A, and disrupted by sonication. Cell lysates were centrifuged at 5000 × *g* for 5 min at 4°C to remove cellular debris. The supernatant was centrifuged at 20,000 × *g* for 1.5 h at 4°C to obtain the pellet, which was solubilized for 1 h in buffer A containing 1% DDM. As those proteins also contains (His)6-sequence in the tag, the membrane proteins were applied to the HisTrap HP affinity column (Cytiva) for purification. The columns were then washed with buffer B (20 mM Tris-HCl, pH 7.5, 150 mM NaCl, 0.05% DDM), followed by wash with buffer B containing 50 mM imidazole. The recombinant proteins were eluted with buffer B containing 500 mM imidazole and concentrated with an Amicon ultra centrifugal filter (10 kDa MWCO, Merck Millipore, UFC501096). The purified recombinant proteins were separated by SDS-PAGE (5%-20%) and further analyzed by staining with Coomassie brilliant blue R-250.

### *In vitro* assay for LLO pyrophosphatase

The Man_5_GlcNAc_2_-DLO substrate was prepared as reported previously (Harada et al., 2013a). Enzyme activity assays were performed in 50 mM MES-NaOH (pH 6.0) containing 200 mM NaCl, 5 mM CaCl_2_, and 0.25% NP-40. Standard LLO pyrophosphatase assay conditions were as follows: Man_5_GlcNAc_2_-DLO (5 pmol) and membrane fractions (10 μg) or purified protein (100 ng) were incubated in a 50 μL scale at 37°C. After the indicated time (for membrane fractions: 16 h; for purified protein: 1 h), the reaction was stopped by adding three volumes of ethanol. After centrifugation at 15,000 × *g* for 5 min, the supernatant was dried by Speed Vac.

Released phosphorylated oligosaccharides were purified as described previously (Harada et al., 2021; Li et al., 2025). Briefly, the dried reaction product was resuspended in 1 mL water and desalted using a PD-midi G-25 column (Cytiva, 17635375). The desalted sample was loaded onto a Sep-Pak Accell QMA (Waters, WAT020545). The QMA column was washed with 10 mL buffer (10 mM Tris-HCl, pH 7.4), and phosphorylated oligosaccharides were eluted with 5 mL elution buffer (10 mM Tris-HCl, pH 7.4, containing 70 mM NaCl). The eluate was divided equally into two samples and incubated with or without 1 μL (1 U) rAPid alkaline phosphatase (Roche, 04898133001) for 16 h at 37°C. The two samples were desalted using a graphitized carbon column (InertSepGC, GL Science, 5010-68000) and dried for the next step of fluorescence labeling.

### Pyridylamination of oligosaccharides (PA-oligosaccharides)

Dried oligosaccharides were labeled with PA (Fujifilm, 011-14181) (Hase, 1994; Hirayama et al., 2010), and excess free PA was removed using a MonoFas I spin column (ANIMOS, A01-1103) as described previously (Hirayama et al., 2010).

### Quantification of PA-oligosaccharides

PA-oligosaccharides were separated by size fractionation HPLC with a Shodex NH2P-50 4E column (4.6 × 250 mm, Shodex, F7630001) as reported previously (Harada et al., 2013a; Hase, 1994). The elution was performed using a gradient of two buffer systems as follows: buffer A, 93% acetonitrile in 0.3% acetate (pH7.0; adjusted with ammonia); buffer B, 20% acetonitrile in 0.3% acetate (pH 7.0). The column temperature was set to 25°C. The gradient program was set at a flow rate of 0.45 mL/min: 0-5 min, 3% buffer B; 5-8 min, 3%-33% buffer B; 8-40 min, and 33%-71% buffer B. PA-oligosaccharides were detected by measuring fluorescence (excitation wavelength, 310 nm; emission wavelength, 380 nm). The amount of PA-oligosaccharides was quantitated using the PA-glucose oligomer (Takara, 4108) as a standard.

### Spot assay of *E. coli* growth

*E. coli* stains were grown in LB at 37°C overnight. 40 μL overnight culture was inoculated into 2 mL LB and incubated at 37°C for 5 h. The OD_600_ was normalized to 1, and serial 10-fold dilutions were made in a 96-well plate. Of each diluted suspension, 5 μL was spotted onto the indicated LB agar media. The growths were observed after 24 h of incubation at 37°C.

### Vancomycin susceptibility analysis

*E. coli* stains were grown in LB at 37°C overnight. Overnight cultures were inoculated into LB medium containing 0.2 mM IPTG with different vancomycin (FUJIFILM Wako, 226-01306) concentrations, with the OD_600_ normalized to 0.01. The OD_600_ of these cultures was measured after growth at 37°C for 6 h (BW25113-WT and *yfiM*-KO) or 16 h (Rosetta-pET28 and pET28-YfiM).

### Measurement of cell lysis by osmotic shock assay

The osmotic shock assay was based on a previous study, with some modifications (Nossal and Heppel, 1966). *E. coli* stains were grown in LB at 37°C overnight. 20 μL overnight culture was inoculated into 2 mL LB and incubated at 37°C until OD_600_ reached 0.6. After adding 0.2 mM IPTG to the culture, the cells were cultured for an additional 3 h and then harvested by centrifugation. After washing once with PBS, the cells were resuspended in either isotonic PBS or pure water, and the ratio of residual optical density (H_2_O/PBS), % OD_600_) was calculated.

### Statistical analyses

The statistical test details are indicated in the figures and figure legends. Statistical analysis was performed by a Student’s t-test. Microsoft Excel was used for statistical analyses.

## Data availability

All data are included in the manuscript and supporting files

## Acknowledgments

Tadashi Suzuki gratefully acknowledges the generous support of late Mr. Kazuhisa Suzuki and Ms. Noriko Suzuki for this research project. This research was supported by the RIEKN Pioneering Project (“Glyco-lipidologue Initiative”), by the Japan Agency for Medical Research and Development-Core Research for Evolutional Science and Technology (AMED-CREST) Grant JP25gm1410006 (to TS).

## Author contributions

ST. L. and Tad. S. designed the experiments. ST. L. performed the experiments. ST. L. and Tad. S. analyzed the results. ST. L. and Tad. S. wrote the manuscript. KP, VL, TN and HT provided bioinformatics information and materials, respectively. All authors reviewed the results and approved the final version of the manuscript.

## Figure Legends

**Figure S1.**
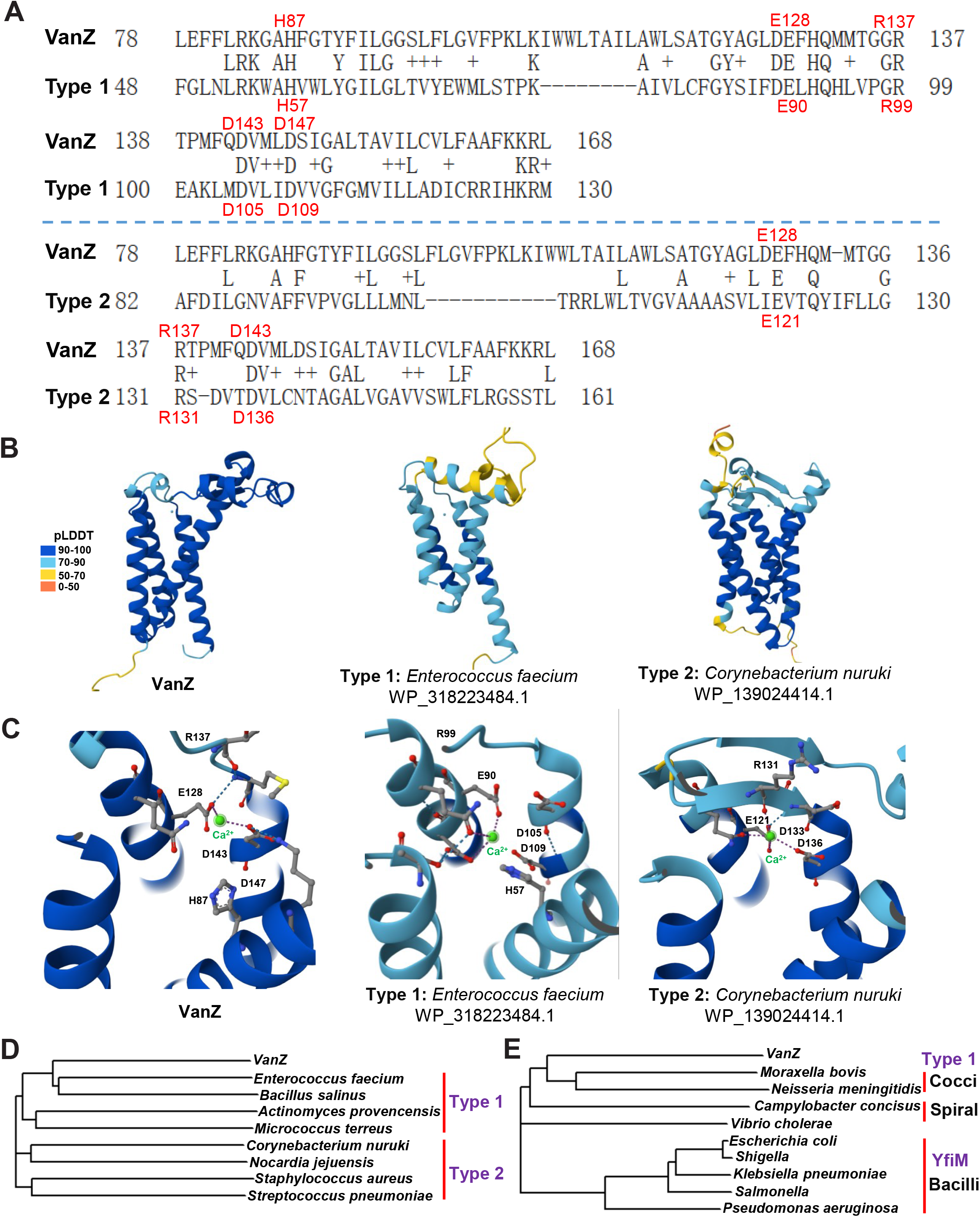
Sequence alignment, structural comparison, and phylogenetic lineages of alternative uncharacterized VanZ/YfiM family proteins. (A) DELTA-BLAST primary sequence alignments of VanZ against uncharacterized Type 1 (top, WP_318223484.1 from *Enterococcus faecium*) and Type 2 (bottom, WP_139024414.1 from *Corynebacterium nuruki*) proteins. Key active site and metal coordinating residues are highlighted and labeled in red (H87, E128, R137, D143, D147 for VanZ; and their corresponding aligned residues in Type 1 and Type 2 proteins). (B) AlphaFold-predicted models for VanZ (left), Type 1 protein (middle), and Type 2 protein (right), colored by the per-residue confidence metric pLDDT on a scale from 0 to 100. The proteins are categorized into Type 1 and Type 2 distinct lineages based on their transmembrane domain (TMD) topologies and helical packaging. (C) Close up view of the predicted catalytic centers. The spatial geometry of critical active site residues coordinating the divalent calcium ion (Ca^2+^, green spheres) remains structurally superimposable among VanZ (left), Type 1 (middle), and Type 2 (right) proteins, demonstrating an invariant tertiary blueprint despite low primary sequence identity. (D, E) Separate phylogenetic trees illustrating the evolutionary segregation and classification of the VanZ/YfiM protein superfamily. Panel D depicts the differentiation between the Gram-positive Type 1 clade (*Enterococcus, Bacillus, Actinomyces, Micrococcus*) and the Type 2 clade (*Corynebacterium, Nocardia, Staphylococcus, Streptococcus*). Panel E details the distinct phylogenetic clustering within Gram-negative bacteria, where the newly identified YfiM clade is strictly restricted to the bacilli lineage (rod-shaped bacteria, including *Escherichia, Shigella, Klebsiella, Salmonella, Pseudomonas*), remaining separate from the Type 1 clusters found in Gram-negative cocci (*Moraxella, Neisseria*) and spiral bacteria (*Campylobacter, Vibrio*).

## References

Abramson, J., J. Adler, J. Dunger, R. Evans, T. Green, A. Pritzel, O. Ronneberger, L. Willmore, A.J. Ballard, and J. Bambrick. 2024. Accurate structure prediction of biomolecular interactions with AlphaFold 3. Nature:1–3.

Aebi, M. 2013. N-linked protein glycosylation in the ER. Biochimica et Biophysica Acta (BBA)-Molecular Cell Research. 1833:2430–2437.

Albalat, R., and C. Cañestro. 2016. Evolution by gene loss. Nature Reviews Genetics. 17:379–391.

Arthur, M., F. Depardieu, C. Molinas, P. Reynolds, and P. Courvalin. 1995. The vanZ gene of Tn1546 from Enterococcus faecium BM4147 confers resistance to teicoplanin. Gene. 154:87–92.

Arthur, M., C. Molinas, F. Depardieu, and P. Courvalin. 1993. Characterization of Tn1546, a Tn3-related transposon conferring glycopeptide resistance by synthesis of depsipeptide peptidoglycan precursors in Enterococcus faecium BM4147. Journal of bacteriology. 175:117–127.

Baba, T., T. Ara, M. Hasegawa, Y. Takai, Y. Okumura, M. Baba, K.A. Datsenko, M. Tomita, B.L. Wanner, and H. Mori. 2006. Construction of Escherichia coli K-12 in-frame, single-gene knockout mutants: the Keio collection. Molecular systems biology. 2:MSB4100050.

Bennett, H.C., and A.M. Mitchell. 2025. Recent advances in understanding of enterobacterial common antigen synthesis and regulation. Open Biology. 15:250055.

Bertani, B., and N. Ruiz. 2018. Function and biogenesis of lipopolysaccharides. Ecosal plus. 8:10.1128/ecosalplus.ESP-0001-2018.

Boratyn, G.M., A.A. Schäffer, R. Agarwala, S.F. Altschul, D.J. Lipman, and T.L. Madden. 2012. Domain enhanced lookup time accelerated BLAST. Biology direct. 7:12.

Brown, S., J.P. Santa Maria Jr, and S. Walker. 2013. Wall teichoic acids of gram-positive bacteria. Annual review of microbiology. 67:313–336.

Burroughs, A.M., G.G. Nicastro, and L. Aravind. 2025. The lipocone superfamily, a unifying theme in metabolism of lipids, peptidoglycan and exopolysaccharides, inter-organismal conflicts and immunity. Elife. 14:RP108061.

Cacan, R., B. Hoflack, and A. Verbert. 1980. Fate of oligosaccharide-lipid intermediates synthesized by resting rat-spleen lymphocytes. European Journal of Biochemistry. 106:473–479.

Cacan, R., A. Lepers, M. Belard, and A. Verbert. 1989. Catabolic pathway of oligosaccharide-diphospho-dolichol: Subcellular sites of the degradation of the oligomannoside moiety. European journal of biochemistry. 185:173–179.

Clark, J.W. 2023. Genome evolution in plants and the origins of innovation. New Phytologist. 240:2204–2209.

Garde, S., P.K. Chodisetti, and M. Reddy. 2021. Peptidoglycan: structure, synthesis, and regulation. EcoSal Plus. 9.

Grabowicz, M., and T.J. Silhavy. 2017. Envelope stress responses: an interconnected safety net. Trends in biochemical sciences. 42:232–242.

Guest, R.L., M.J. Lee, W. Wang, and T.J. Silhavy. 2023. A periplasmic phospholipase that maintains outer membrane lipid asymmetry in Pseudomonas aeruginosa. Proceedings of the National Academy of Sciences. 120:e2302546120.

Harada, Y., R. Buser, E.M. Ngwa, H. Hirayama, M. Aebi, and T. Suzuki. 2013a. Eukaryotic oligosaccharyltransferase generates free oligosaccharides during N-glycosylation. J Biol Chem. 288:32673–32684.

Harada, Y., C. Huang, S. Yamaki, N. Dohmae, and T. Suzuki. 2016. Non-lysosomal degradation of singly phosphorylated oligosaccharides initiated by the action of a cytosolic endo-β-N-acetylglucosaminidase. Journal of Biological Chemistry. 291:8048–8058.

Harada, Y., K. Nakajima, S. Li, T. Suzuki, and N. Taniguchi. 2021. Protocol for analyzing the biosynthesis and degradation of N-glycan precursors in mammalian cells. STAR protocols. 2:100316.

Harada, Y., K. Nakajima, Y. Masahara-Negishi, H.H. Freeze, T. Angata, N. Taniguchi, and T. Suzuki. 2013b. Metabolically programmed quality control system for dolichol-linked oligosaccharides. Proceedings of the National Academy of Sciences. 110:19366–19371.

Hase, S. 1994. High-performance liquid chromatography of pyridylaminated saccharides. Methods Enzymol. 230:225–237.

Hase, S., T. Ikenaka, and Y. Matsushima. 1979. Analyses of Oligosaccharides by Tagging the Reducing End with a Fluorescent Compound I. Application to Glycoproteins. The Journal of biochemistry. 85:989–994.

Hirayama, H., J. Seino, T. Kitajima, Y. Jigami, and T. Suzuki. 2010. Free oligosaccharides to monitor glycoprotein endoplasmic reticulum-associated degradation in Saccharomyces cerevisiae. J Biol Chem. 285:12390–12404.

Hsu, A.-F., J.W. Baynes, and E.C. Heath. 1974. The role of a dolichol-oligosaccharide as an intermediate in glycoprotein biosynthesis. Proceedings of the National Academy of Sciences. 71:2391–2395.

Jeong, H., V. Barbe, C.H. Lee, D. Vallenet, D.S. Yu, S.-H. Choi, A. Couloux, S.-W. Lee, S.H. Yoon, and L. Cattolico. 2009. Genome sequences of Escherichia coli B strains REL606 and BL21 (DE3). Journal of molecular biology. 394:644–652.

Kawakami, N., and S. Fujisaki. 2018. Undecaprenyl phosphate metabolism in Gram-negative and Gram-positive bacteria. Bioscience, Biotechnology, and Biochemistry. 82:940–946.

Leive, L. 1965. Release of lipopolysaccharide by EDTA treatment of E., coli. Biochemical and biophysical research communications. 21:290–296.

Li, S.-T., K. Kamada, A. Honda, J. Seino, T. Matsuda, T. Suzuki, N. Dohmae, Y. Shichino, S. Iwasaki, and Y. Noda. 2025. LLP1 is a pyrophosphatase involved in homeostasis/quality control of dolichol-linked oligosaccharide. Journal of Cell Biology. 224:e202501239.

Manat, G., S. Roure, R. Auger, A. Bouhss, H. Barreteau, D. Mengin-Lecreulx, and T. Touzé. 2014. Deciphering the metabolism of undecaprenyl-phosphate: the bacterial cell-wall unit carrier at the membrane frontier. Microbial Drug Resistance. 20:199–214.

Massarweh, A., M. Bosco, S. Iatmanen-Harbi, C. Tessier, N. Auberger, I. Chantret, C. Gravier-Pelletier, and S.E. Moore. 2016. Demonstration of an oligosaccharide-diphosphodolichol diphosphatase activity whose subcellular localization is different than those of dolichyl-phosphate-dependent enzymes of the dolichol cycle. Journal of Lipid Research. 57:1029–1042.

Nagaraja, T.G. 2022. Basic bacteriology. Veterinary microbiology:11–28.

Nikaido, H. 2003. Molecular basis of bacterial outer membrane permeability revisited. Microbiology and molecular biology reviews. 67:593–656.

Nossal, N.G., and L.A. Heppel. 1966. The release of enzymes by osmotic shock from Escherichia coli in exponential phase. Journal of Biological Chemistry. 241:3055–3062.

Pavlova, M., A. Asaturova, and A. Kozitsyn. 2022. Bacterial cell shape: Some features of ultrastructure, evolution, and ecology. Biology Bulletin Reviews. 12:254–265.

Pootoolal, J., J. Neu, and G.D. Wright. 2002. Glycopeptide antibiotic resistance. Annual review of pharmacology and toxicology. 42:381–408.

Roosild, T.P., J. Greenwald, M. Vega, S. Castronovo, R. Riek, and S. Choe. 2005. NMR structure of Mistic, a membrane-integrating protein for membrane protein expression. Science. 307:1317–1321.

Saha, S., S.R. Lach, and A. Konovalova. 2021. Homeostasis of the Gram-negative cell envelope. Current opinion in microbiology. 61:99–106.

Schjoldager, K.T., Y. Narimatsu, H.J. Joshi, and H. Clausen. 2020. Global view of human protein glycosylation pathways and functions. Nature reviews Molecular cell biology. 21:729–749.

Schoch, C.L., S. Ciufo, M. Domrachev, C.L. Hotton, S. Kannan, R. Khovanskaya, D. Leipe, R. Mcveigh, K. O’Neill, and B. Robbertse. 2020. NCBI Taxonomy: a comprehensive update on curation, resources and tools. Database. 2020:baaa062.

Scott, P.M., K.M. Erickson, and J.M. Troutman. 2019. Identification of the functional roles of six key proteins in the biosynthesis of Enterobacteriaceae colanic acid. Biochemistry. 58:1818–1830.

Silhavy, T.J., D. Kahne, and S. Walker. 2010. The bacterial cell envelope. Cold Spring Harbor perspectives in biology. 2:a000414.

Strasser, R. 2016. Plant protein glycosylation. Glycobiology. 26:926–939.

Studier, F.W., P. Daegelen, R.E. Lenski, S. Maslov, and J.F. Kim. 2009. Understanding the differences between genome sequences of Escherichia coli B strains REL606 and BL21 (DE3) and comparison of the E. coli B and K-12 genomes. Journal of molecular biology. 394:653–680.

Yamamoto, N., K. Nakahigashi, T. Nakamichi, M. Yoshino, Y. Takai, Y. Touda, A. Furubayashi, S. Kinjyo, H. Dose, and M. Hasegawa. 2009. Update on the Keio collection of Escherichia coli single-gene deletion mutants. Molecular systems biology. 5:335.

Zhang, Y., and J. Skolnick. 2005. TM-align: a protein structure alignment algorithm based on the TM-score. Nucleic acids research. 33:2302–2309.

